# Thermodynamic uncertainty relation optimizes copy number of signaling proteins

**DOI:** 10.1101/2025.01.04.631284

**Authors:** Aryan Dubey, Shreyansh Verma, Bhaswar Ghosh

**Affiliations:** Center for Computational Natural Sciences and Bioinformatics, International Institute of Information Technology, Hyderabad, India

## Abstract

Cell signaling systems are involved in sensing changes in the environment by activating a set of transcription factors (TF) that typically diffuse within the nucleus to trigger transcription of the required genes. The TFs can diffuse randomly in and out of the nucleus leading to fluctuations in different components which severely limits the accuracy of estimating environmental input. The diffusion usually happens through the nuclear pore complexes which enables the tf protein to enter the nucleus either passively or actively depending on it’s size. In this study, we explored the role of diffusion on the information transmission capacity of a set of tfs using a coupled mathematical and machine learning approaches to experimental data in yeast under several stress conditions. We found that the the activation followed by biased diffusion of transcription factors (TF) towards the nucleus triggers amplifying magnitude of the overall TF currents towards the nucleus as well as reduces the fluctuations. In fact, to our surprise the diffusion rate estimated from the data is found to be positively correlated with the protein mass, indicating the possibility of active diffusion since a negative correlation is expected in case of passive diffusion. The active diffusion in fact facilitates faster entry to the nucleus enabling faster information transmission with nuclear protein concentration as output. Additionally, higher copy number of the TF also improves information transmission by reducing overall noise in the output. However, improved information owing to faster active diffusion and higher copy number comes at an extra cost of increased entropy production due to the inherent thermodynamic uncertainty relation (TUR)for non-equilibrium systems. A linear optimization analysis demonstrates a corelation between the protein size and the optimized protein number which corroborates the actual observation. Thus, experimental measurements coupled with diffusion based theoretical models demonstrate the role of diffusion on optimizing cellular information processing.

## INTRODUCTION

Every living cell is required to sense the environment properly to ensure appropriate phenotypic response. The sensing of the extracellular signal is usually carried out by a receptor on the cell membrane which then triggers a downstream signaling cascade finally leading to the activation of one or more transcription factors. These activated transcription factors then diffuse inside the nucleus to stimulate transcription of several genes which would enable the cell to alter its phenotype appropriate for the environmental signal [1]. The biochemical pathways starting from the signal transduction to the nuclear localization involve inherently stochastic processes giving rise to fluctuation in the transcription factor output. These are the basis for noisy response stemming from both intrinsic and extrinsic sources and finally severely limiting the accuracy of estimating the input signal which may lead to inappropriate activation of the downstream genes [2, 3, 4, 5]. Consequently, undesired activation of the genes would not only impact the phenotypic outcome as well as incur unnecessary waste of the cell’s valuable resources in terms of material and energy. Thus, accurate estimation of the environmental signal is crucial for the cell to avoid such undesirable outcomes. cells use several strategies in order to improve the accuracy of estimation, namely negative feedback to reduce the fluctuation, amplifying the output to increase the signal to noise ratio or multiple measurement of the signal. Recent studies illustrated that one of the ways the multiple measurements can be performed is through dynamic measurement the output at multiple time points which is able to improve the information transmission capacity significantly provided the output is dominated by extrinsic noise. The role of extrinsic and intrinsic sources of fluctuations had been one of the topics of several studies in the last two decades both theoretically and experimentally in different contexts of biological systems including gene expression and cellular signaling. For example, the extrinsic noise was shown to be the dominating factor in signaling pathway in yeast where the extrinsic noise emerges from the cell to cell variability of the pathway protein numbers. Most of these previous studies have concentrated on the noise arising from the molecule numbers and consequently the experiments were designed to decouple intrinsic and extrinsic sources stemming from the same which essentially measured the protein number by fluorescent tagged proteins at the single cell level. But, these techniques did not capture the noise arising from diffusion of the protein molecules in and out of the nucleus which would contribute to enhancing the intrinsic noise. In the last few years with the unprecedented advancement of microfluidic devices coupled to high resolution imaging techniques, it is now possible to track localization of individual transcription factors inside a single cell. Recently, an experimental study has been conducted to capture the localization of a set of transcription factors in yeast S. Cerevisiae under multiple different environmental stress and information transmission capacity of the pathways [6]. These experimental techniques can effectively be harnessed to quantify the noise arising from the diffusion process. However, how the diffusion process and its associated fluctuation would impact the signaling accuracy is not well explored.

In this study, we take recourse to these publicly available data sets and first employ information theory techniques to calculate the accuracy of information transmission carried by the nuclear localization of the activated proteins. The change in the output (e.g. stimulated expression of a gene) carries information about the input variable. In general, the precision with which the input value can be estimated from measuring the output improves with larger changes (i.e. dynamic range) of the output and with lower output noise. According to Cramėr–Rao inequality, the error in estimating the input from the output is bounded by the Fisher information, defined as the relative entropy change of the output distribution for an infinitesimal change in the input around a given input value [7]. Furthermore, information transmission capacity of signalling systems can also be estimated by calculating the mutual information, which is increasingly being used to characterize biochemical signalling networks [8, 9, 10, 11, 12, 13, 14]. The mutual information measures the mutual interdependence between the input and the output distributions by calculating the relative entropy of the output distributions conditioned on the input with respect to the unconditioned output distributions. In both cases, more information can be extracted about the input from the output distribution if the relative entropy change is large [15, 16, 17].

As the experiments measure the nuclear to cytoplasmic ratio of the TFs over time, the time traces encode the diffusion dynamics of the TFs from cytoplasmic to nucleus and consequently the diffusion rate can be estimated by calculating the fluctuation time scale of the time traces. The diffusion time scale essentially captures the memory of the dynamics in time. One of the intriguing issue which naturally arises from the analysis is to identify the factors that determine the variable diffusion rates among the TFs. We expect that in case of passive diffusion the the diffusion rate would reduce with mass, but we observed a positive correlation between the mass and the diffusion rate indicating the possibility of active diffusion. Passive macromolecular diffusion through nuclear pore complexes (NPCs) is thought to decrease dramatically beyond a 30–60 kDa mass threshold [18]. In fact, all the TFs in our case have mass more than 40KDa which is naturally points towards the need for active diffusion process and the positive correlation stemms from the increased number of active sites [19] inside the Nuclear Localisation Signal (NLS) sequences as the length of the sequence increases.

However, it is also generally established that improving accuracy in input estimation will in turn impose extra burden by enhancing energy dissipation [20, 21, 22, 23]. In fact, the general principle of thermodynamic uncertainty relation (TUR) postulates the need for higher entropy production to improve accuracy by reducing fluctuation in the current in a random walk scenario [24]. The nuclear localization of transcription factors under the influence of environmental stress offers a conducive scenario to explore the principle of TUR in biological context. Previous studies have already alluded to TUR in biology specifically in the context of molecular motors [25]. But, it’s application in signal transduction cascades remains unexplored plausibly due to lack of experimental data. In the last few years with the unprecedented advancement of microfluidic devices coupled to high resolution imaging techniques, it is now possible to track localization of individual transcription factors inside a single cell. Recently, an experimental study has been conducted to capture the localization of a set of transcription factors in yeast S. Cerevisiae under multiple different environmental stress and information transmission capacity of the pathways [6].

In addition to the inherent stochasticity, the underlying reactions involved in biochemical pathways are essentially out of equilibrium processes. The close connection between information and heat dissipation in non-equilibrium processes is established on a strong theoretical footing owing to the resolution of the long standing puzzle of Maxwell’s demon through the Landauer erasure principle in the last century [26, 27]. Entropy production rate is the measure of heat dissipation in non-equilibrium processes and entropy production rate also represents the energy consumption in many systems at steady state [28]. The essential phenomena which separates a non-equilibrium process from an equilibrium one is the presence of currents arising from breaking of time reversal symmetry [29, 30, 31]. The entropy production rate can be exactly quantified from the the trajectory of a stochastic processes by calculating current if the underlying equation giving rise to the dynamics is known [32, 33], But, unfortunately the underlying dynamical equations are not known in majority of the cases. One of the alternative strategies is to determine the current fluctuation from the thermodynamic force by training a neural network model based on trajectory data sets and subsequently estimate a bound on the entropy production rate [34]. Another technique relies on the time reversal symmetry breaking criteria to estimate entropy production rate [35]. Here, we used these methods to estimate the entropy production rates of the trajectories of protein localization along with the calculation of accuracy of information transmission mentioned previously. The information transmission capacity is observed to display a strong correlation with the estimated bound on the entropy production rate expected from the thermodynamic uncertainty relation. In fact, a model based on biased random walk of the proteins fundamentally corroborates the analysis of experimental data. The estimated diffusion rates from the trajectories exhibit a positive correlation with masses of TFs as expected from the general principle of active diffusion process and the overall fluctuation of the trajectories reduces with protein copy number. Thus, the information transmission capacity also improves with mass and protein copy number. The entropy production on the other hand would also amplify with mass and protein copy number posing a trade-off scenario which essentially confers a constraints on the optimized protein copy number depending on its mass. Our study offers a general theoretical framework to lay down an underlying mathematical principle of TUR which drives the protein localization through energy dissipation and optimality in information transmission.

## RESULTS

### Reduction in fluctuation due to amplified response

To explore the role of fluctuation on the response of the biological signaling pathway, we analyzed the single cell measurement of the dynamics of the transcription factor nuclear localization signal under various stresses in the yeast *S. Cerevisiae* [6]. In this experiment (Appendix E), the nuclear localization signal of ten different transcription factors under three different stresses, namely carbon, oxidative, and osmotic stresses, is estimated using fluorescent tagged transcription factors through a microfluidic devise. The nuclear localization is quantified by the ratio of the fluorescent signal within the nucleus to the cytoplasm. The experimental data provide us with the trajectories before and after the stress inside individual cells for each transcription factors (Fig S1). The cell to cell variability in nuclear localization after the treatment of the stress for one of the transcription factors quantifies the noise in the response for that particular transcription factor. The average signal provides a general trend in response of the transcription factors (Fig 1a). To estimate the relationship between the average amplitude and fluctuation, we calculated the mean response at the maximum magnitude for each transcription factor (Fig S1) after the stress is applied and the corresponding variance describes the noise strength. The noise represented by the coefficient of variation 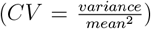 shows an expected decreasing trend with the maximum average response for the threes different stresses (Fig 1b-1c). Thus, as the response is higher the noise in the response is lower. The lower noise as a result would improve the signal-to-noise ratio, leading to an improvement in information transmission.

**FIG. 1:**
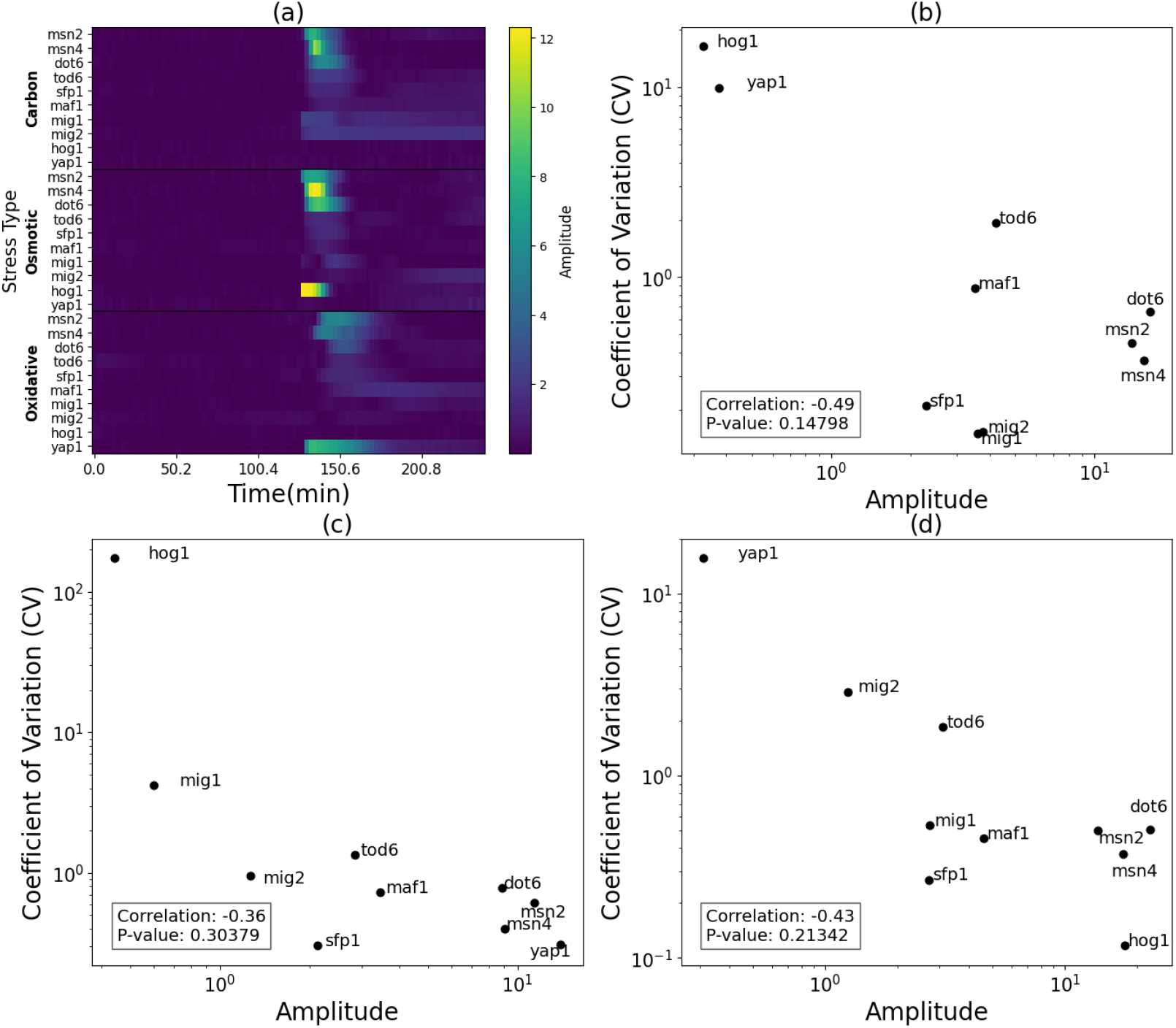
High response amplitude reduces fluctuation. (a)The heat map shows the average trajectories of the amplitude of nuclear-cytoplasmic ratio (NC ratio) over 200 cells for different transcription factors before and after application of the stress under different stress conditions as indicated by an increase in the NC ratio just after the stress is applied.(b) The reduction in fluctuation as the response amplitude is higher, described by the relationship between the 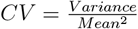 of the maximum amplitude for individual single cell trajectories after stress with its *Mean* calculated over 200 cells for carbon stress (c) the same calculation is done for osmotic stress and (c) oxidative stress

### Improved information transmission due to reduced fluctuation

Information transmission can be quantified by calculating the mutual information between input and output. Mutual information provides a measure of information the output signal carries about the change in the input. As in our case, there are two values of the stresses (eg. in presence and absence of carbon stress) which entails only one bit of information at the input and thus, we would expect the output would carry at most 1 bit of information about the input. However, some information is going to be inevitably lost due to fluctuation arising from stochasticity in the underlying pathway. We used a previously described method to calculate the information [6]. The method utilizes a SVM based classifier to predict the input from the output signal at a particular time point or the entire trajectory (Methods section D). The confusion matrix provides the joint distribution *p*(*X* = *x, Y* = *y*) of the predicted value x and actual value y which can be used to determine the mutual information between the two variables *X* and *Y* following the definition

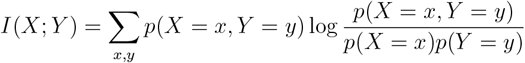

where *p*(*X* = *x*) and *p*(*Y* = *y*) represent the total probability distribution of *X* and *Y* respectively given by

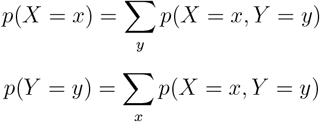

Using this method, the mutual information is calculated at each time point along trajectory using a sliding time window as indicated in Fig 2a-c for different stress conditions and select the mutual information value where the response is maximum for each transcription factors. The mutual information indeed increases as the response is higher irrespective of the nature of the stress which corroborates our hypothesis that the higher response improves information transmission by reducing the fluctuation for carbon (Fig 2d), osmotic (Fig 2e) and oxidative stresses (Fig 2f) as depicted by correlations (R=0.67,p=0.03 for carbon; R=0.68,p=0.001 for osmotic; R=0.92,p=0.0002 for oxidative) between response amplitude and information.

**FIG. 2:**
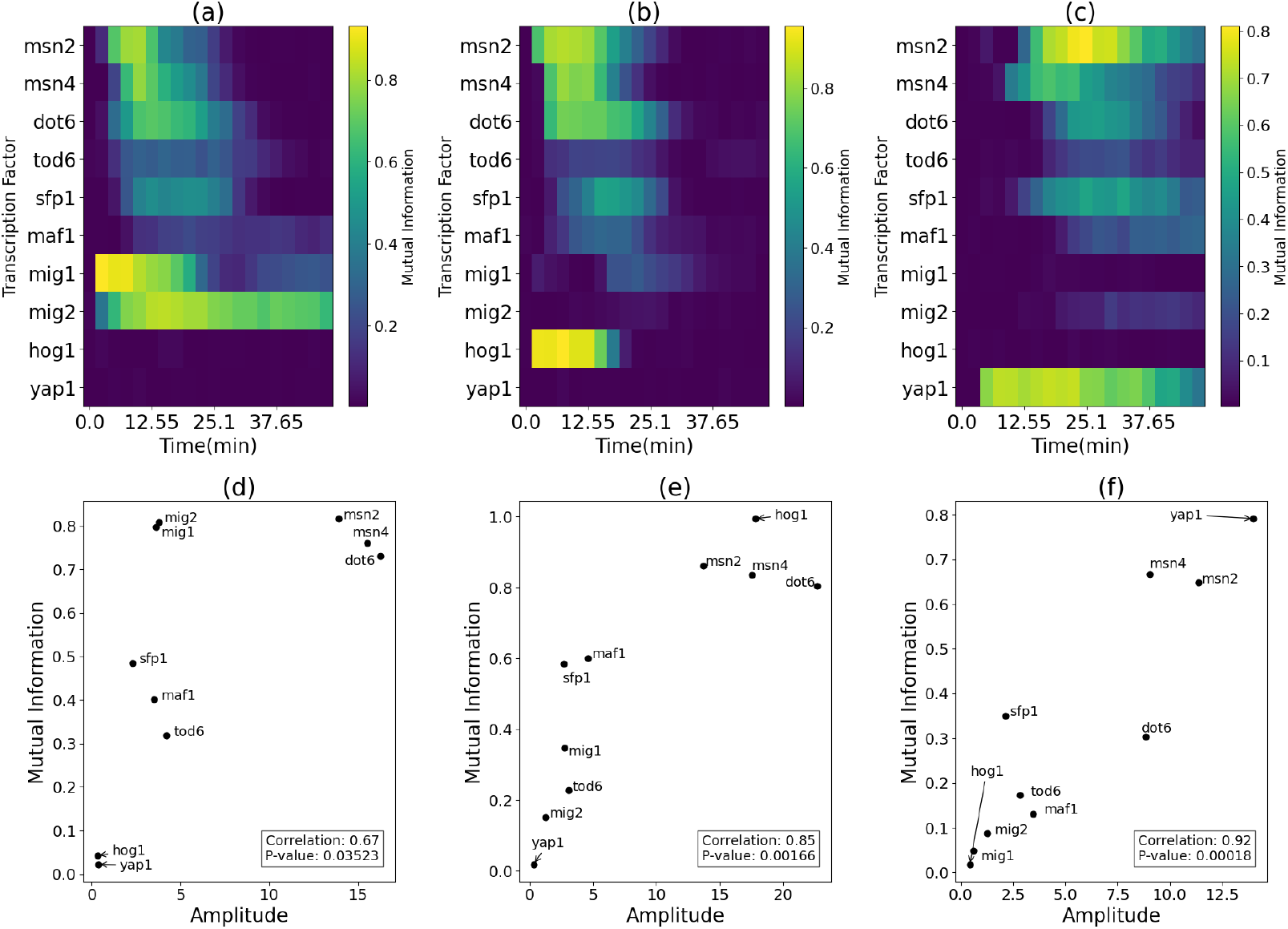
Information transmission improves for higher response amplitude. The mutual information was calculated at the maximum response amplitude for each transcription factor with stress values as the input. (a) The trajectories of mutual information with time for different transcription factors calculated by taking a sliding time window along the response trajectories for carbon stress (b) osmotic stress (c) oxidative stress. The mutual information at maximum response increases with the average response amplitude for the three stress condition namely (d) Carbon (e) Osmotic and (f) Oxidative stress indicated by the positive correlation between mutual information and response amplitude.

### Higher response enhances the entropy production rate

The thermodynamic uncertainty relation [36] is described by the inequality

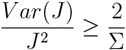

where J represents the current through the physical system and Σ corresponds to the entropy production in the system due to the current. The uncertainty relation essentially states that the reduction in fluctuation entails a cost by increasing the entropy production. As in our case, reduction in fluctuation due to amplified response must incur higher entropy production in the nuclear localization process. We take recourse to one of the recently developed techniques [34] to estimate the entropy production rate from the trajectory of the nuclear localization signal from the current fluctuation determined by training a neural network model(Appendix C).Following the statement of uncertainty relation, the entropy production rates would be higher at higher response amplitude since fluctuation is lower, leading to the observed correlation between the mutual information and entropy production for the transcription factors over carbon (Fig. 3a,R=0.81), osmotic (Fig. 3b,R=0.56) and oxidative stresses (Fig. 3c,R=0.74). The shaded regions represent 90 percent confidence level from regression models. The ratio between the mutual information (I) and *log*(1 + Σ) is defined as an efficiency of each TF and we observed that the efficiency is always less than 1 (Fig. 3d-3f). In fact, while a 50 to 100 fold variability in *log*(1 + Σ) or mutual information exists among the TFs, only around five fold variability in efficiency is observed due to the presence of the correlation.

**FIG. 3:**
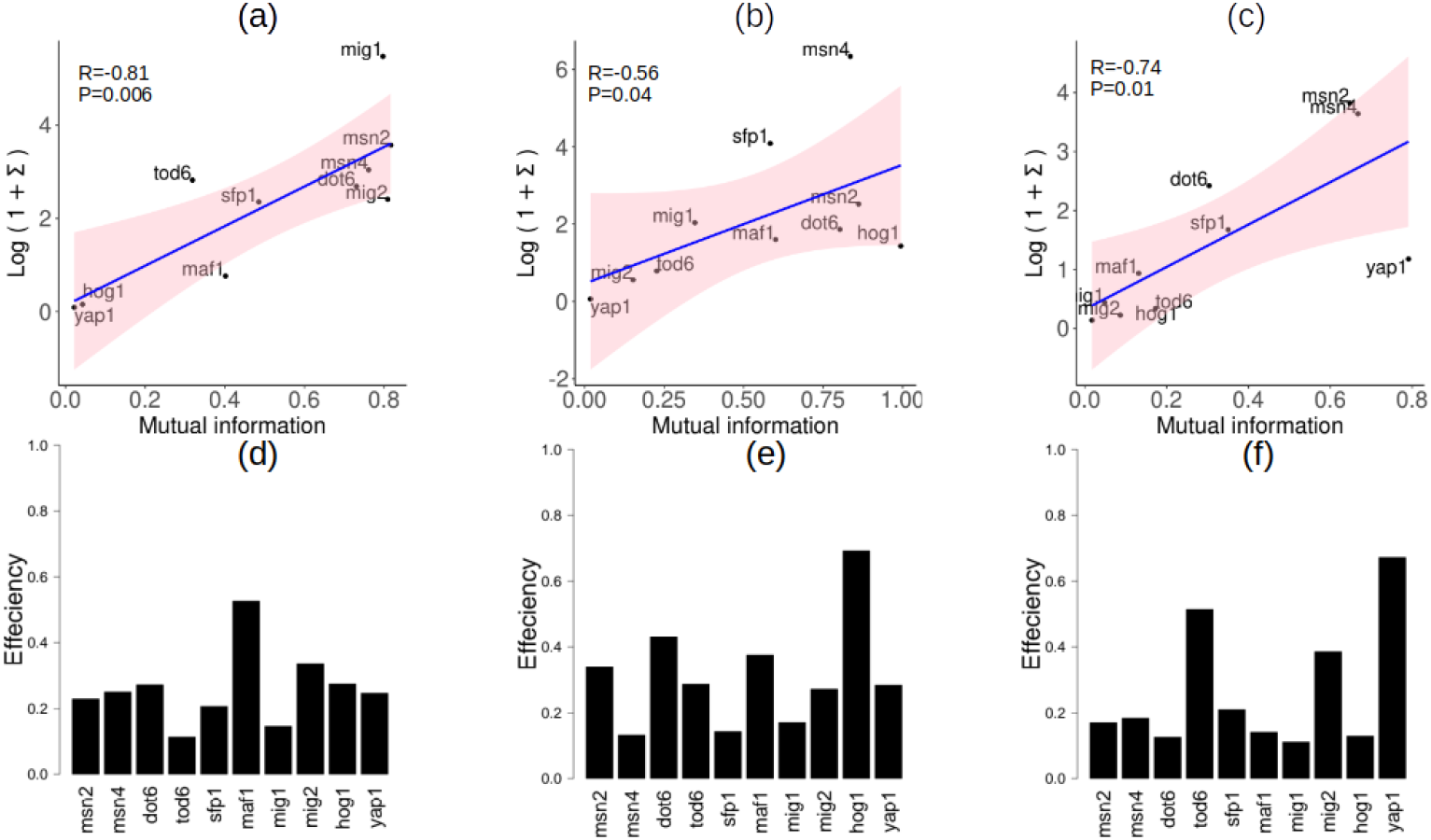
Entropy production limits information transmission through the signaling pathways. The thermodynamic uncertainty relation sets an upper bound on mutual information *I < log*(1 + Σ). The entropy production rate for each trajectory was calculated using a neural network model as described in the main text. The average entropy production over the time to reach the maximum amplitude is determined for each transcription factor at the three different stress conditions. The positive correlation between *log*(1 + Σ) and mutual information for (a) carbon (b) osmotic (c) oxidative stresses suggest the requirement for higher entropy production in reducing the fluctuation leading to higher mutual information. The values of *log*(1 + Σ) is always greater then the mutual information values as indicated by the values of the efficiencies for (a)carbon (b) osmotic and (c) oxidative stresses for the TFs.

### Entropy production constraints the information transmission by imposing an upper bound

When the environmental input has two possible values, The mutual information between the input and output can be derived as (Appendix A)

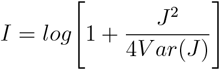

where J is the flux through the pathway as defined in the previous section. In view of the uncertainty relation, the information would have an upper bound given by the inequality

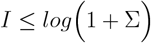

In fact, this inequality assumes that the time scale of the change in the input and the output are slow so that the pathway relaxes to the steady state along with the input. The estimated bounds from the entropy production rates are indeed higher than the mutual information in our case as indicated in the previous figure 2. The algorithm based on neural network as described in the previews section relies on the time reversal symmetry-breaking criteria to estimate the entropy production rate. This method essentially calculates the forward and reverse transition probabilities between a set of preselected states. The selection of the states are later optimized to minimize the entropy production rate using optimization algorithms. In fact, the estimated bound in this method is found to be higher than the mutual information, although since it calculates the entropy production also for low flux the correlation between flux and the entropy production rate was not very high. In this method the entropy production rate is calculated from the trajectory of the experiment which essentially measures the ratio between the nuclear and cytoplasmic protein. Further, the experimental data does not provide us the flux and it’s fluctuation through the pathway. The true entropy production as well as the fluctuation in the current can only be calculated if the underlying reaction network is known. The actual estimation of entropy production is shown in a previous study.

### Active diffusion rate is proportional to the protein mass

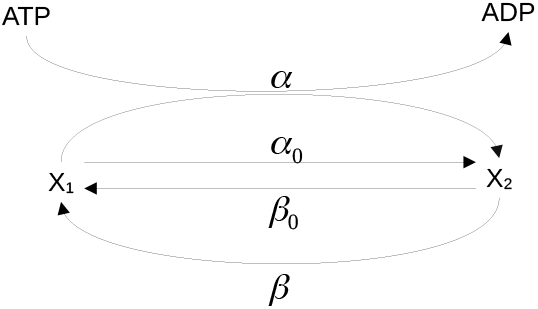

In order to obtain a technique to estimate the diffusion rate from the NC ratio measurement, we first constructed simple mathematical model of diffusion as described in the above figure where each TF molecule can actively diffuse in the nucleus at rates *α* and passively at a rate *α*_0_. It can also diffus out of the nucleus at a rate *β* actively and at a rate *β*_0_ passively. If *x*_1_ and *x*_2_ are the concentration of the TF inside the cytoplasm and nucleus respectively then the dynamics of *x*_2_ will be governed by the differential equation,

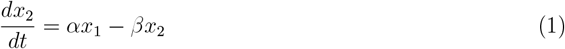

Here, we clubbed the two rates *α* and *α*_0_ together as *α* = *α* + *α*_0_, the two rates *β* and *β*_0_ together as *β* = *β* + *β*_0_ If, the ratio 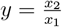 and *x*_1_ + *x*_2_ = *X*_*T*_, the equation for y is given by,

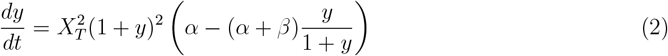

The steady state of *y* is given by 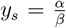 and linearizing the dynamics around steady state and adding white noise, one obtains,

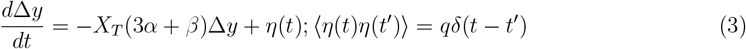

Before the application of stress the NC ratio is one, i.e, *α* = *β*. The covariance can be written as

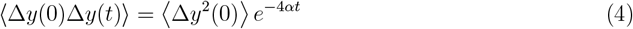

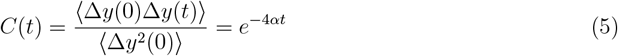

Which implies that measuring the autocorrelation of the dynamics of y would provide us with an estimate of the diffusion rate *α*. We calculate the diffusion rate *α* by calculating autocorrelation *C*(*t*) at different time points from the trajectories for individual TF and taking the slope of the *C*(*t*) vs *t* line in the log-scale (Appendix F). The diffusion rates are found to be linearly correlated with the mass of the protein (extracted from SGD database) which indicates an active diffusion process through the nuclear pore (Figure 4a), since for equilibrium diffusion, the diffusion rate is expected to vary inversely with mass.

**FIG. 4:**
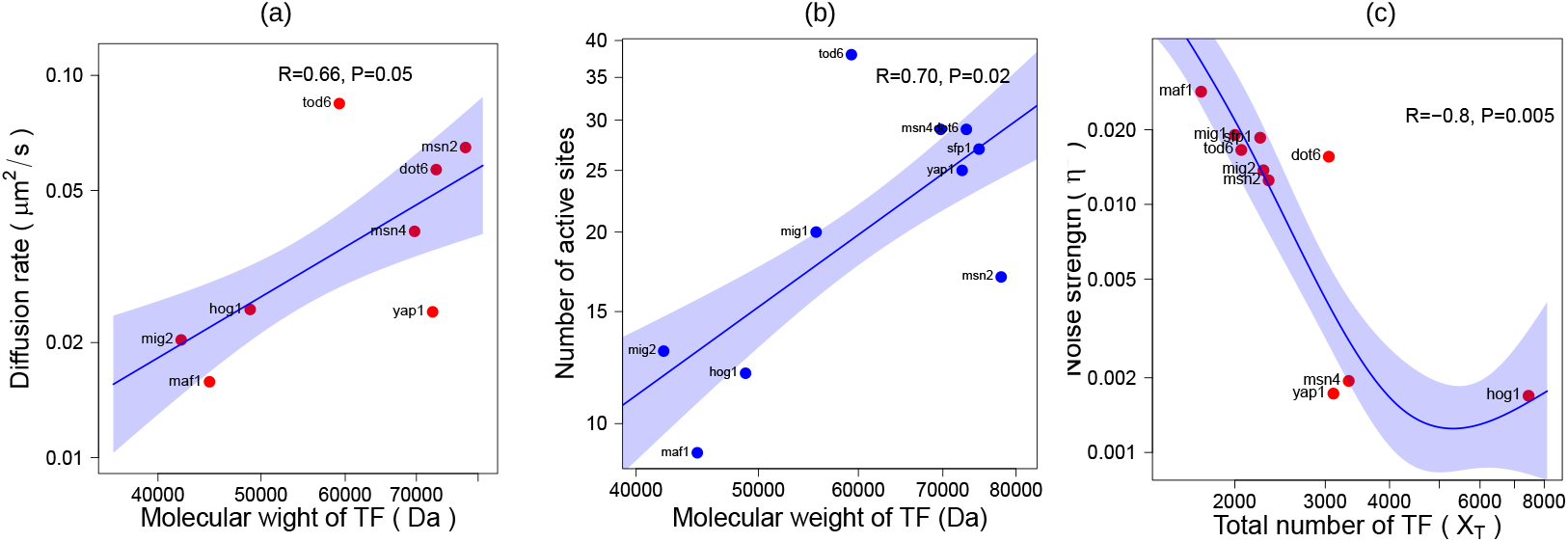
Information improves with mass and protein copy number. (a)The diffusion rate of nuclear localization estimated from the trajectories are positively correlated with the protein masses as indicated. The blue shadow represents the 90 percent confidence interval of a regression model. (b) The number of active sites for Nuclear Localisation Signals (NLS) as a function of the mass of the protein. The blue shadow represents the 90 percent confidence interval of a regression model. (c) The coefficient of variation (*η*^2^ = *CV*) for unstressed trajectories. The blue shadow represents the 90 percent confidence interval of a spline fitting model.

In order to investigate the plausible reason of the relationship between the diffusion rate and protein mass, we utilized a a recently developed a highly accurate NLS prediction program (cNLS Mapper [19]), which calculates NLS activity scores by using activity-based, but not sequence-based, profiles for different classes of importin-*α*-dependent NLSs. To assess the functional contribution of various amino acids at every position in each of 3 monopartite NLS classes, all of the amino acid residues of the template NLSs, as representatives of these classes, were serially replaced with 20 other amino acid residues. The relative nuclear import activities of these altered sequences were assayed in yeast and ranked in 1 of the 10 levels based on the localization phenotype of the GFP reporter. From the cNLS mapper output, the total number of active sites (*N*_*s*_) which influences the nuclear import was calculated and the *N*_*s*_ values were found to be correlated with mass, i.e, the length of the protein (Figure 4b). This probably indicates that the higher length of a protein enhances the chance of having more active NLS residues triggering diffusion through nuclear pore complex.

### Noise reduces with protein copy number

The value of the variance in *y*, i.e, ⟨Δ*y*^2^⟩ be calculated as below,

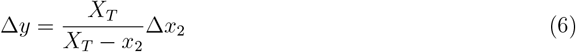

And from this, one can substitute the values of 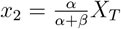 and 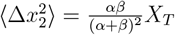 at steady state to obtain,

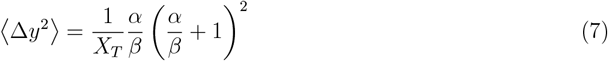

which at unstressed condition (*α* = *β*) turns out to be,

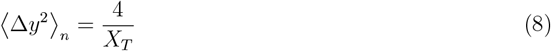

This implies that the variance in the unstressed condition would provide an estimate of the total number TF. In fact, the overall CV value is observed to be systematically reduced with the protein number (Figure 4c).

### Information improves with both protein copy number and mass

As indicated in Equation(8) the fluctuation in the NC ratio before stress depends on the number of the TF, molecule. In order to compare the relative fluctuation of different TFs after stress, it is required to normalize the fluctuation of the response before stress. To achieve the normalization, we defined the normalized response *R* which guarantees the mean and variance to be 0 and 1 respectively before the treatment of stress irrespective of the TF, as

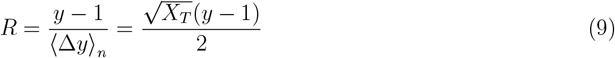

which implies that, 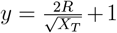. Thus, following the equation(7) where 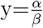. To estimate the signal to noise ratio (SNR), we assume that the stressed and unstressed conditions have equal probability to calculate the conditional variance, 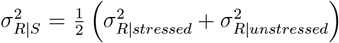. Since, *R* = 0 for unstressed condition,

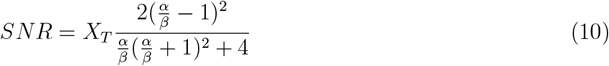

The mutual information can be written as,

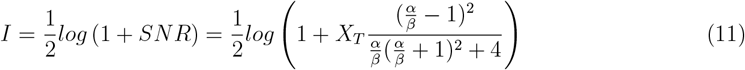

which implies

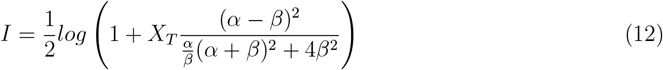

The forwards and reverse diffusion rate would be proportional to the mass of the protein in case of active diffusion. The forward diffusion rate would be triggered by the stress Thus, *α* = *K*_1_*m* + *α*_0_ and *β* = *K*_2_*m* + *β*_0_. Thus, we can write (Appendix B)

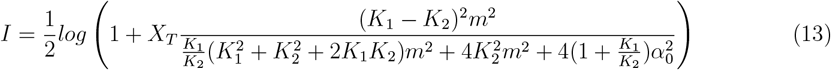

which implies that the information increases as the protein number *X*_*T*_ increases by reducing the overall fluctuation and the further improvement happened due to enhanced active diffusion rate as mass increases. However, the increase in protein number and mass are also supposed to amplify the entropy production.

### Breaking of detailed balance and entropy production

For our case, the cytoplasmic version *x*_1_ and nuclear version *x*_2_ of the transcription factor would follow the dynamics,

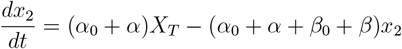

For the chemical reaction system described above, the transitions between the two states are driven by an external force due to nuclear import if the TF by the enzyme at a rate *α* and a nuclear export at a rate *β*. The nuclear import is triggered by the active diffusion process requiring energy of GTP hydrolysis in addition to the passive diffusion driven equilibrium transition rates *α*_0_ and *β*_0_. External driving breaks the detailed balance of the cycle, leading to the entropy production and corresponding power consumption by the cycle [37]. The steady state is given by

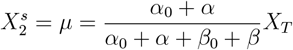

The corresponding entropy production rate at steady state is,

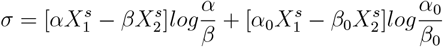

At steady state, the fluxes would be equal

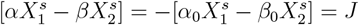

This essentially breaks the detail balance as the fluxes are not individually zero which is responsible for the net fluxes and the emergence of a nonzero entropy production rate at steady state. Thus, finally the expression for the flux is given by,

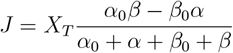

The entropy production rate is given by

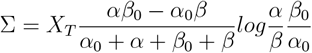

The external driving force exerted by the enzyme *α* associate a free energy F to the reaction in addition to the equilibrium free energy. Thus one can associate the reaction rates with the driving force free energy and the equilibrium reaction rates [38],

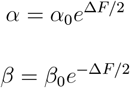

which implies that

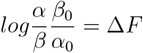

Additionally, if the free energy is zero at equilibrium,

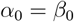

With this assumption, the entropy production rate or the power consumption at steady state is finally takes the form,

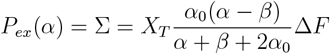

Here, Δ*F* corresponds to the free energy change of ATP hydrolysis. Thus, entropy would increase as the output increases.

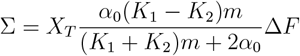

### A trade-off optimizes protein copy number

Since both the information and the corresponding entropy production increases with mass *m* and protein number *X*_*T*_, following the TUR, we can define a fitness function *L* considering the information transmission as benefit and the entropy production as a cost.

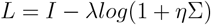

assuming Lagrange multiplier *λ*, so that *L >* 0 and *η* represents a proportional factor for the effect of entropy production on the reduction in cell growth. Thus, one can write,

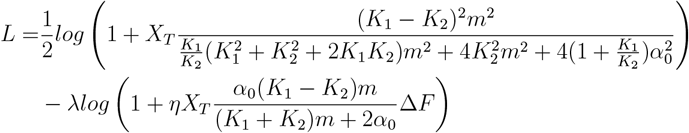

To calculate the optimal value of *X*_*T*_ for a particular value of mass *m*, 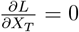 which implies,

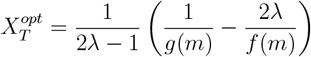

where, 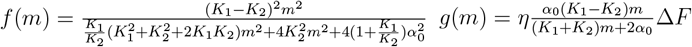

The optimum 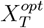 changes with the mass of the TF. The condition for the case when 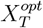 increases with *m*, i.e., 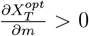, is given by,

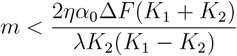

Thus, till an upper bound on mass is reached, the optimal protein copy number 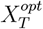 would increase with *m*, but it will start to decrease as the mass crosses the upper bound. It is notable that the the upper bound on mass is high when the ratio 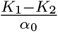 is low. In fact, the fitness *L* increases at low values of *X*_*T*_ and reaches a maximum and reduces subsequently (Figure 5a) at different values of mass *m*. The optimum value of 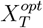 where the maximum fitness is attained increases with *m* (Figure 5b red line) when (*K*_1_−*K*_2_) values are low. However, for higher (*K*_1_−*K*_2_) values, at maximum fitness (Figure 5c), the optimum 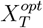 vales tend to reduce with *m*. (Figure 5b blue line). Interestingly, this pattern is observed in the actual relation between mass and protein number (Figure 5d) where for a set of TFs (mig1, mig2, sfp1, tod6 and maf1) the protein number increases with mass probably (blue line in Figure5d)corresponding to low responses and for another set (msn2, msn4, dot6, yap1 and hog1), the protein number reduces with mass (red line in Figure 5d), representing higher response. This result indicate that the protein number may have been optimized by the cell depending on its mass following the TUR based fitness described above. In fact, the set of proteins belonging to the first set exhibits much lower response than the second set of proteins compared over the multiple stresses considered here corroborating the hypothesis of optimality (Figure 5e). Response at different doses of the stress also show similar trend (Figure 5f).

**FIG. 5:**
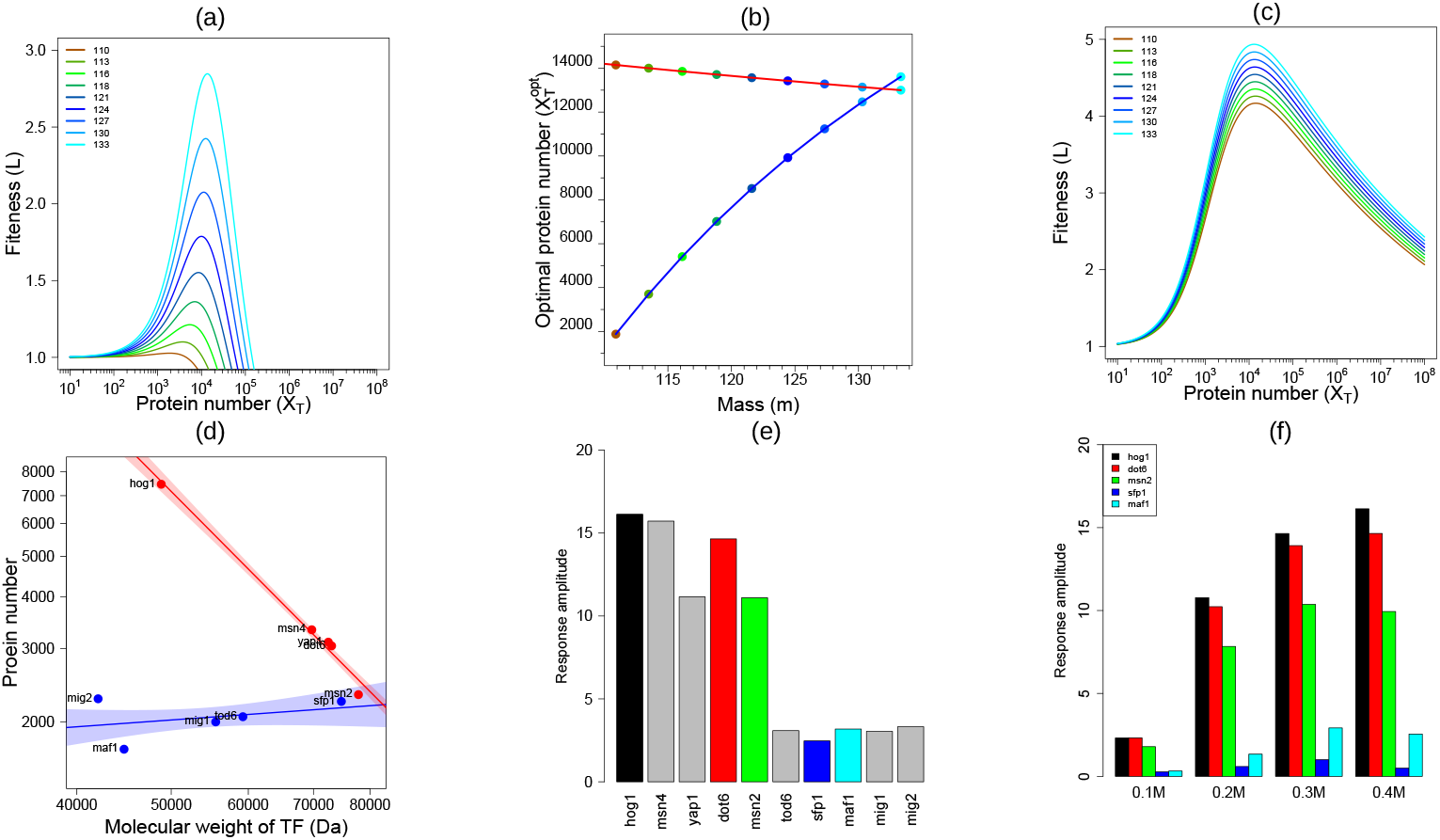
Protein copy number is optimized by an Information vs entropy production trade-off. The thermodynamic uncertainty relation sets an upper bound on mutual information *I < log*(1 + Σ). The fitness(*L*) is defined as *L* = *I λlog*(1 + Σ). (a) The values of *L* as a function of protein copy number (*X*_*T*_) at different values of protein mass *m* as indicated for *K*_1_ = 1.1; *K*_2_ = 1.0; *λ* = 1.09; *η* = 1; *α*_0_ = .3; Δ*F* = 0.01 where 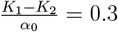 is small. (b) The optimal values of *X*_*T*_ where *L* is maximum as a function of mass *m* (red line). (c) The values of *L* as a function of protein copy number (*X*_*T*_) at different values of protein mass *m* as indicated for *K*_1_ = 1.3; *K*_2_ = 1.0; *λ* = 1.09; *η* = 1; *α*_0_ = .1; Δ*F* = 0.01 where 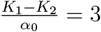 is large. The red line in reprsnets the functional relation between the optimal *X*_*T*_ and mass *m* in this case. (d) The relationship between the protein copy number and mass for the TFs in consideration. (e) The comparison of response amplitudes for the TFs over all the three stresses. (f) The comparison of response amplitudes for osmotic stresses at different levels as indicated.

## DISCUSSION

The requirement for optimal resource allocation due to the energy-accuracy trade-off in signaling systems is emerging as a crucial design principle for biological signaling systems in multiple studies [39, 20, 40, 41, 42]. Disparate studies investigated different aspects of the biological systems to present energetic trade-offs scenarios. The role of energetic cost in cellular signaling systems has been implicated in previous studies [39, 22, 21, 20]. Apart from cellular signaling, the similar consideration in shaping the rates in reaction networks was investigated for error proof reading in translation [43], as well as in copying biochemical systems [44]. in the context of bacterial chemotaxis where the past memory remains stored in the slow methylation patterns in the receptor where as the phosphorylation signal relaxes back to pre-treatment state [45]. This was achieved through the utilization of an integral negative feedback. In fact, the studies also reinforce the fact that the memory storage through the negative feedback imposed extra energetic cost and cell anchors the feedback loop in order to optimize the energy-accuracy trade-off in scenarios of energy budget constraints [21]. The evolutionary tuning of protein copy number or proteome resource allocation were demonstrated in previous studies by cost-benefit analysis [46, 42]. Despite several such studies primarily dealing with optimality principle shaping the design of biological systems, a concrete theoretical foundation which can integrate different studies into a coherent mathematical framework is missing. Thermodynamic uncertainty relation (TUR) can offer such an underlying framework to fill the gap and lay down a foundational ground for uncovering general principles driving architecture of biological systems through evolutionary landscape.

TUR essentially establishes connection between the fluctuation and entropy production in far from thermal equilibrium system. This formulation in fact deals with the long standing observation of Maxwell’s demon which alluded to a fundamental connection between information and thermodynamics [47]. According to Landauer’s principle [26], the work done during a thermodynamic cycle occurs due to erasure of memory The thermodynamic uncertainty relation essentially reinforce the connection by constraining the fluctuation in a physical current and power consumption in the system and finally fluctuation’s intimate link with information establishes the role of power consumption to information as discussed in our study specifically in the context biological signaling as an example of a physical system [48, 49]. Derivation of the basic uncertainty relation presupposes some assumption for the system namely, stationarity and linear response [50]. In the case we studied the stationarity condition only holds true at the beginning of the application of stress but the pathway activity starts to fall down after some time. In order to establish the validity of the results found from the experimental data, a mathematical model which essentially takes into account the stationarity condition illustrated that the overall claim is not contingent on the non-stationarity at the later part of the pathway activity. In fact, the pathway activity almost becomes very low i.e, close to the equilibrium state after some time which indicates that current as well as entropy production becomes vanishing small in that regime,

Our study exhibits the limit to the information transmission in view of the energy cost due to dissipation at steady state and living organism may deal with this trade-off in multiple ways either by reducing the ATP consumption in signaling cascade which will increase fluctuation degrading information transmission, or introducing feedback loop or storing past information as memory while shutting down the pathway output. Irrespective of the strategy, TUR presents an upper bound on the information given the entropy production by the pathway. In this study, we explored the information carried by a set of TFs actively diffusing through the the nuclear membrane and the corresponding entropy production. The mass of the protein plays a crucial role in import of the active TF through the nuclear pores under stress as a bigger protein would contain more number of active nuclear port sites. The protein copy number plays a role by reducing overall fluctuation in the nuclear localization signal. As a result, on one hand, both the mass and protein number tend to improve information transmission, but on the other hand, impose an extra cost by increased entropy production, posing a trade-off scenario for the cell to resolve. We argued that the cell deal with the situation by optimizing the protein copy number and the optimized protein number 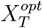 systematically changes with the mass of the protein depending on the immediate response after stress. Our analysis reveal that 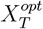 values increases with mass when response is low and reduces with mass when response is high. It indicates that when response is low, the cell could not afford to produce higher protein number for reducing fluctuation to improve information, whereas the sinuation is different when response under stress is high. There can be several contributing factors which sets the response amplitude apart from the diffusion rate and protein number, e.g. activation rate of the TFs through phosphorylation.

In the paper from which we have taken the data[6], the transcription factors were selected based on generalists and specific types. Some transcription factors are found to be generalists which are activated under all the three stresses and some few which are only activated under specific stresses. Generalists (Msn2/4, Tod6 and Dot6, Maf1, and Sfp1) can encode the nature of multiple stresses, but only if stress is high; specialists (Hog1, Yap1, and Mig1/2) encode one particular stress, but do so more quickly and for a wider range of magnitudes. The aim of our paper however is to demonstrate that irrespective of the stress or transcription factor, the information or accuracy of signaling is limited by the entropy production to provide a general biophysical perspective on the operation of the signaling. Moreover, the experimental data is not stationary since the cells start to adapt after some time under all stresses, only adaptive dynamics are stress specific. However, the main aim of the model is to estimate the reaction rates which will capture the maximum response which happens very fast and takes a much longer time scale (~ 15 minutes) to adapt compared to the reaction time scales in seconds. Of course it is a quasi steady state assumption at the maximum response to capture the entropy production and information at maximum response. In the mathematical model, we assumed that the total number of TFs remains the same before and after the stress which ignores any transcriptional regulation neglecting the entropy production due to protean production.

## ACKNOWLEDGMENTS

Authors thank Department of Biotechnology (No. BT/RLF/Re-entry/32/2017), Government of India for funding this project.

## APPENDIX

### A. Bound on mutual information

If *P* (*R, S*) is the joint distribution of the response R and signal S, total variance of the response [11]

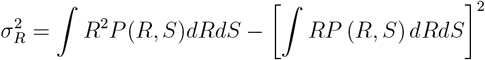

Conditional Variance

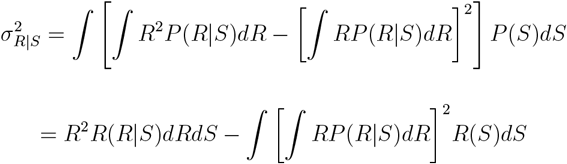

Thus,

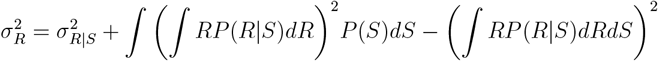

If there are two inputs *S*_1_ and *S*_2_

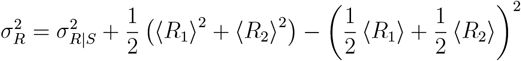

Where,

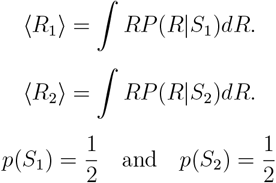

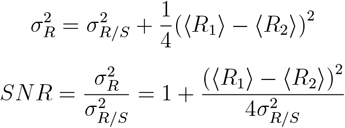

If we assume that the distributions are Gaussian.

The mutual information

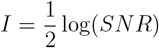

If the response is taken as the current through system.

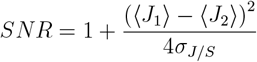

We assume that the system is at equilibrium when no stress is applied and it goes out of equilibrium when stress is applied.

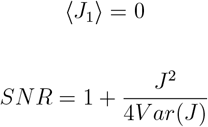

According to the thermodynamic uncertainty relation

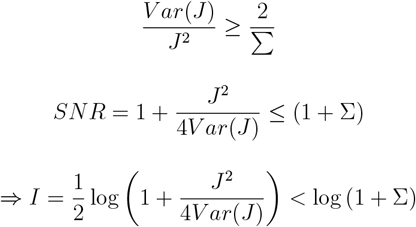

#### B. Signal to Noise ratio for the model under active diffusion

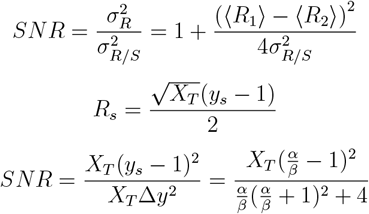

From the figure, *α* = *α* + *α*_0_, *β* = *β* + *β*_0_ and *α*_0_ = *β*_0_ at equilibrium. Thus, we can write,

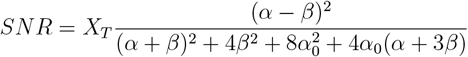

The forwards and reverse diffusion rate would be proportional to the mass of the protein in case of active diffusion. The forward diffusion rate would be triggered by the stress Thus, *α* = *K*_1_*m* and *β* = *K*_2_*m*. Thus, we can write,

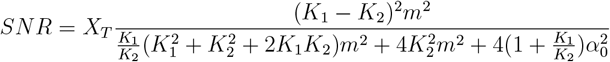

We assumed that, *α* and *β* being larger than *α*_0_, the linear terms are ignored.

#### C. Estimating Mutual Information

The dataset used in the study consists of nuclear localization (reported as the ratio of nuclear fluorescence to cytoplasmic fluorescence) of different transcription factor in presence of different stress values. The data is sampled at an interval of 2.5 minutes[6]. The 10 transcription factors used in the study are namely:

**msn2, msn4, dot6, tod6, sfp1, maf1, mig1, mig2, hog1, yap1**

The three types of stress in which their dynamics is studies are namely.

##### Carbon Stress, Osmotic Stress, Oxidative Stress

We followed a methodology similar to the one outlined in the referenced paper [6]. The data is normalized by taking the mean and variance of the pre-stress points. Specifically, we consider a trajectory of length 40, comprising 20 pre-stress points and 20 post-stress points, with the point of stress applied at the midpoint of the trajectory. The mean *µ* and variance *σ* are calculated from the 20 pre-stress points, ensuring that the pre-stress mean is 0 and the pre-stress variance is 1.

##### 1. Mutual Information through Decoding

To calculate Information Gain, we employed a decoding-based technique similar to the one described in the referenced paper [**cite**]. The Information Gain *I* is defined as:

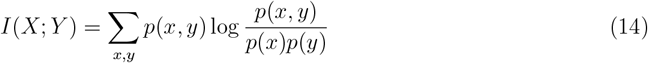

Direct evaluation of this equation is challenging due to the high dimensionality of *p*(*x, y*). Instead, we utilize a parametric form of *p*(*x, y*).

The decoding-based estimator operates on individual trajectories, determining whether each trajectory corresponds to a control or stress condition. The classifier functions as a deterministic mapping *f* : *x* → ŷ that assigns a trajectory *x* to an environment state ŷ.

The classifier’s performance is summarized in a confusion matrix, which shows the counts of correctly and incorrectly classified instances:

| | | Predicted ($\hat{s}$) | |
| --- | --- | --- | --- |
| | | $\hat{s}_0$ | $\hat{s}_1$ |
| Actual ($s$) | $s_0$ | $P(\hat{s}_0, s_0)$ | $P(\hat{s}_1, s_0)$ |
| | $s_1$ | $P(\hat{s}_0, s_1)$ | $P(\hat{s}_1, s_1)$ |

This matrix is used to estimate the joint probabilities necessary for calculating mutual information. The entries in the matrix are normalized to obtain joint and marginal probabilities, which are then used in the mutual information formula.

The steps of the algorithm are as follows:

##### 1. Normalize the Data

For each time point *x*_*i*_(*t*), compute the normalized value 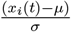. This normalization is based on the first 20 time points before stress is applied and the 20 time points after stress is applied, totaling 40 time points.

##### 2. Split the Dataset

Divide the dataset into two halves: the first half as **control** and the second half as **stress**.

##### 3. Train-Test Split

Further split the dataset into training (70%) and testing (30%) subsets. Use a **Support Vector Machine (SVM)** to classify the trajectories.

##### 4. Generate Confusion Matrix

Post-classification, generate a confusion matrix that shows the count of correct and incorrect classifications.

##### 5. Normalize Confusion Matrix

Normalize the confusion matrix to create a probability distribution.

##### 6. Calculate Information Gain

Compute the Information Gain *I* using the normalized probabilities in the mutual information formula given in equation 14

This approach provides an approximate value for the Information Gain using parametric conditional probabilities.

#### D. Estimated EPR using a neural network model

##### 1. Data Preparation

To prepare the data, we followed a procedure similar to that used for Mutual Information analysis. The data was normalized by computing the mean and variance of the pre-stress points. Specifically, we considered trajectories of length 40, comprising 20 pre-stress points and 20 post-stress points.

##### 2. Algorithm

In a previous study [34], a novel algorithm is presented for calculating the Entropy Production Rate (EPR) from time series data of biological systems using machine learning techniques. We adopted the framework outlined in this study to calculate the EPR for the translocation of transcription factors between the cytoplasm and nucleus. This framework provides the EPR directly from the trajectory.

After obtaining a time series of EPR values, we computed their mean to obtain a single EPR value for each transcription factor. The entropy production estimation framework (NEEP) employs a feedforward neural network (FNNKt in the non-stationary case) because of its flexibility in approximating nonlinear force fields. We used 2 hidden layers with 30 units each, a learning rate of 10^−3^, Adam optimizer, and 50,000 iterations, selecting the best model by validation score on a held-out test set. To assess robustness, the training is repeated with multiple seeds, the data is perturbed with noise, the trajectory is down sampled and the length is truncated; the entropy production estimates remained stable across these conditions. Finally, interpretability is ensured by visualizing the learned thermodynamic force fields as vector plots and analyzing sensitivities (Jacobian and simplified parametric fits), which connect the neural network outputs to biologically meaningful dynamics. More details regarding train vs test errors are presented in Fig. S2. We utilized the (NEEP) representation along with the Variational Estimator, considering them as the default option for handling unknown cases.

#### E. Experimental method

We briefly describe the protocol of data generation used in the previously published experiment [6], the data of which was used for the entire analysis of the the study.

##### 1. Microscopy and microfluidics

Microscopy and microfluidics 1.2.1 Cell preparation and loading ALCATRAS Overnight cultures were inoculated such that cells would reach mid-log phase by the following morning. Cells were diluted in fresh medium to OD600 0.1 and incubated an additional 3–4 hours for loading into microfluidics devices at OD600 0.3–0.4. To expose multiple strains to the same environmental conditions and to optimize data acquisition, a multi-chamber version of ALCATRAS8, 9 was used, which allowed five different strains to be loaded into distinct chambers but still be exposed to the same extracellular media.

##### 2. Changing the extracellular environments

For the stress-type experiments, medium either containing 0.1 percent glucose (carbon stress) was supplied, supplemented with 0.4 M sodium chloride (osmotic stress), or supplemented with 0.5 mM hydrogen peroxide (oxidative stress). The dye Cy5 was added to the syringe containing stress medium to monitor its arrival time and the sharpness of the switch.

##### 3. Image acquisition and analysis

During each experiment, bright-field and fluorescence images at five z-sections spaced 0.6 micron apart were acquired. The maximum projection of these images (the maximum pixel values across all z-sections) was used for quantification. The Cy5 channel was acquired in a single focal plane. Cells were segmented from bright-field images using the DISCO algorithm 10, 11, which identifies cell centers with a support vector machine (SVM) and cell edges with an active-contour-based method applied to the bright-field z sections. the nuclear accumulation of transcription factors was quantified by calculating the ratio between the average of the five brightest pixels within the cell and the median fluorescence of the whole cell.

## SUPPLEMENTARY MATERIAL

**Fig. S1:**
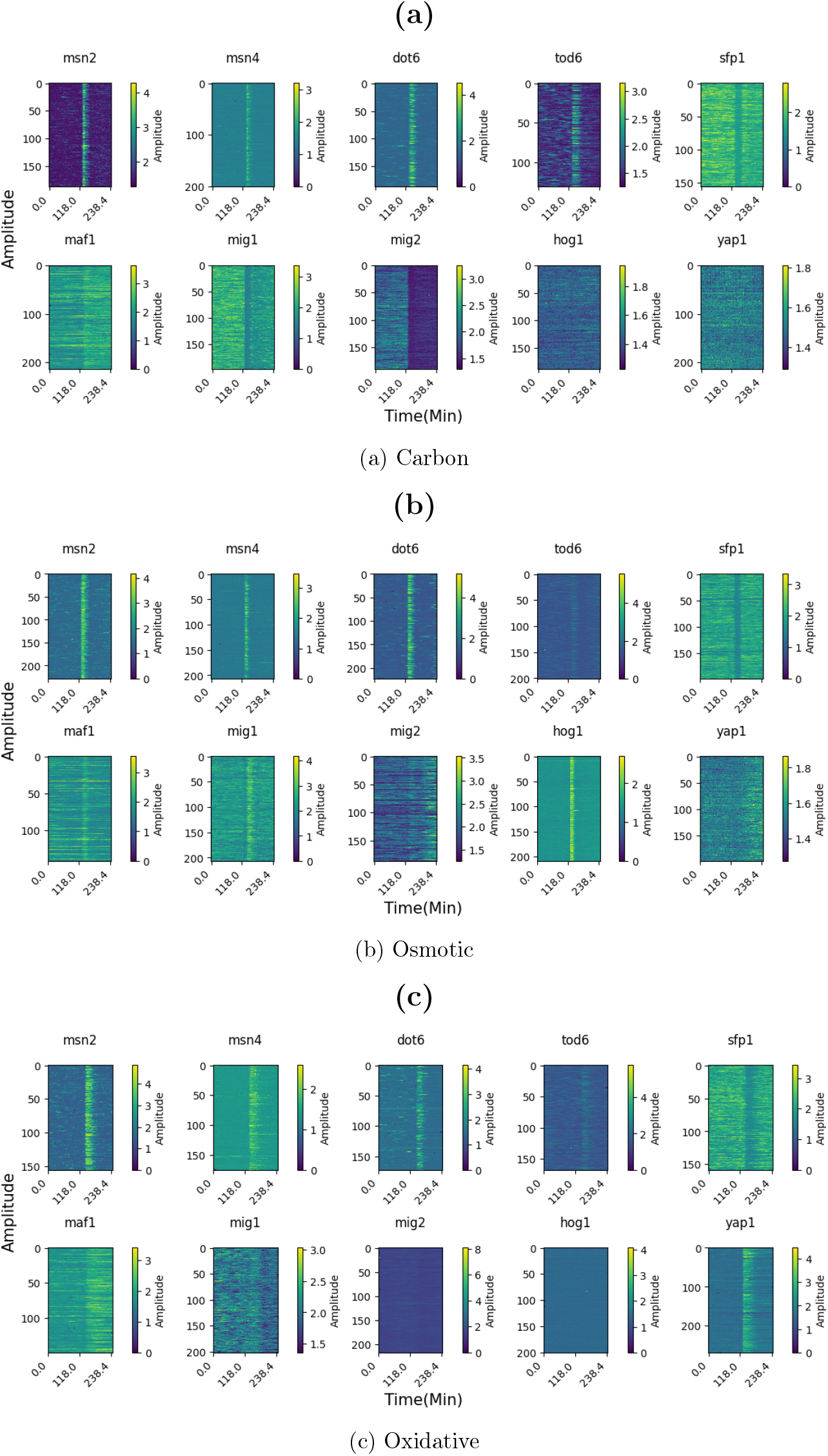
Trajectories of NC ratio inside individual cells for different transcription factors before and after different stress conditions: (a) Carbon, (b) Osmotic, and (c) Oxidative.

**Fig. S2:**
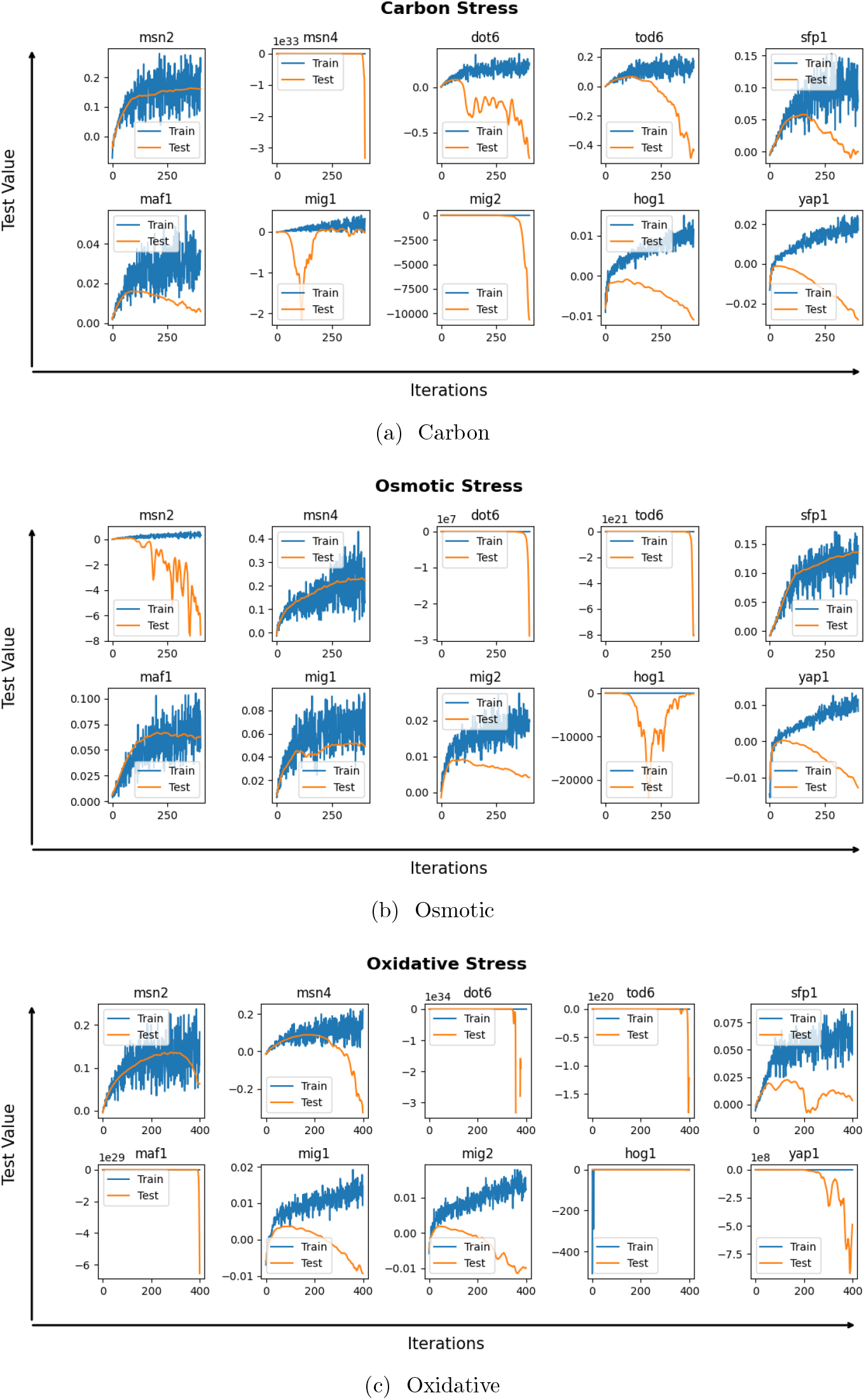
Train vs Test errors for NEEP framework in case of different stress values: (a) Carbon, (b) Osmotic, and (c) Oxidative.

